# Receptor-like role for PQLC2 amino acid transporter in the lysosomal sensing of cationic amino acids

**DOI:** 10.1101/2020.07.15.204800

**Authors:** Gabriel Talaia, Joseph Amick, Shawn M. Ferguson

## Abstract

PQLC2, a lysosomal cationic amino acid transporter, also serves as a sensor that responds to scarcity of its substrates by recruiting a protein complex comprised of C9orf72, SMCR8 and WDR41 to the surface of lysosomes. This protein complex controls multiple aspects of lysosome function. Although it is known that this response to changes in cationic amino acid availability depends on an interaction between PQLC2 and WDR41, the underlying mechanism for the regulated interaction is not known. In this study, we establish that the WDR41-PQLC2 interaction is mediated by a short peptide motif in a flexible loop that extends from the WDR41 β-propeller and inserts into a cavity presented by the inward-facing conformation of PQLC2. This data supports a transceptor model wherein conformational changes in PQLC2 related to substrate transport regulate the availability of the WDR41 binding site on PQLC2 and mediate recruitment of the WDR41-SMCR8-C9orf72 complex to the surface of lysosomes.

## INTRODUCTION

Lysosomes have long been known as sites where hydrolytic enzymes break down pathogens and cellular macromolecules into the basic building blocks of cells (amino acids, nucleic acids, sugars, lipids and metals) so that diverse transporters can return these nutrients to the cytoplasm to support ongoing anabolic needs (1). More recently, it has been established that in addition to clearing potentially toxic material and recycling nutrients, important signals are transduced from lysosomes. This prominently includes signalling via mTORC1 which is initiated on the cytoplasmic surface of lysosomes under the control of a highly regulated network of proteins that sense and respond to changes in nutrient and growth factor availability (1–3). Cellular responses to various pathogens are also initiated from lysosomes via several members of the Toll-like receptor (TLR) family (4). Both mTORC1 and TLR signalling from lysosomes depend on mechanisms to transduce signals from the lysosome lumen to the cytoplasm.

In addition to these well established pathways for signalling from lysosomes, the lysosomal cationic amino acid transporter PQLC2/SLC66A1 was recently found to recruit a heterotrimeric complex comprised of the C9ORF72, SMCR8 and WDR41 proteins to the cytoplasmic surface of lysosomes (5). Knock out studies in multiple model organisms have established that this protein complex is critical for normal lysosome function and has impacts on both mTORC1 and TLR signalling (6–11). Considerable attention has been focused on the C9orf72 gene due to a hexanucleotide expansion in a non-coding region of this gene which causes amyotrophic lateral sclerosis and frontotemporal dementia (12–14). Multiple mechanisms have been proposed to explain how the C9orf72 hexanucleotide expansion causes neurodegenerative diseases and although the expansion occurs in a non-coding region, epigenetic silencing of the affected allele results in an overall decrease in C9orf72 expression levels in carriers of the repeat expansion that has been proposed to have disease relevance (12, 14–19). Understanding PQLC2-dependent recruitment of the C9orf72-SMCR8-WDR41 complex to lysosomes is thus broadly relevant for both lysosome cell biology as well as neurodegenerative diseases that potentially arise from C9orf72 deficiency.

Within the C9orf72-SMCR8-WDR41 protein complex, WDR41 forms a bridge between C9orf72-SMCR8 and PQLC2 and is thus essential for lysosome recruitment of the C9orf72-SMCR8-WDR41 complex (Amick et al., 2020, 2018). This interaction between WDR41 and PQLC2 is negatively regulated by the availability of the cationic amino acids that are transported out of lysosomes by PQLC2 (5). As a result, the C9orf72-SMCR8-WDR41 complex is maximally recruited to lysosomes when cells are starved of cationic amino acids (5, 11, 20). Like other members of the SWEET family of transporters, PQLC2 contains 7 transmembrane spanning segments that are critical for its transporter activity (21, 22). However, PQLC2 contains only minimal sequences that are exposed to the cytoplasm and has no folded cytoplasmic domains as candidates for mediating either amino acid sensing or WDR41 interactions. Therefore, although the PQLC2-WDR41 interaction is central to lysosomal nutrient sensing, it is unknown how PQLC2 acts as both an amino acid transporter and a platform for the regulated recruitment of the C9orf72 complex to the surface of lysosomes.

In this study, we establish the basis for interactions between WDR41 and PQLC2 through an iterative process of structural predictions and biochemical validation. Our results show that a short peptide motif within a flexible loop that extends from the WDR41 β-propeller inserts into the large cavity exposed by the inward facing conformation of PQLC2 recruits WDR41 to lysosomes. Our results thus provide a model for how PQLC2 acts as both a cationic amino acid transporter and WDR41 receptor in order to support the ability of cells to sense and respond to changing cationic amino acid availability.

## RESULTS AND DISCUSSION

### Prediction of a novel mechanism for the WDR41-PQLC2 interaction

At endogenous expression levels, the interactions between WDR41 and PQLC2 are tightly regulated by cationic amino acid availability (5). Defining how this PQLC2-WDR41 interaction controls WDR41-SMCR8-C9orf72 complex recruitment to the surface of lysosomes is important for understanding lysosome-based nutrient sensing and signalling. To establish the mechanistic basis for this regulated interaction, we first performed homology modelling using HHpred (MPI Bioinformatics Toolkit; Zimmermann et al., 2018) to predict a WDR41 structure. The human ODA16 β-propeller protein (PDB: 5NNZ) displayed the highest predicted structural homology with WDR41 and was selected as the template for further WDR41 modelling. This led to a model of WDR41 as an 8-bladed β-propeller with a prominent loop (or unstructured region) that is formed at the bottom of blade 7, between β-strands C and D (7CD loop; Fig. 1A). The secondary structure predicted by WDSP, a database for WD40 proteins, also predicted a similar large loop extending from the WDR41 β-propeller (Ma et al., 2019; Y. Wang et al., 2015). Although not available at the start of our study, two recently reported cryo-EM structures for the WDR41-C9orf72-SMCR8 complex validate this prediction for the overall structural organization of WDR41 and showed that the interaction with SMCR8-C9orf72 is via the WDR41 β-propeller (23, 24).

**Figure 1:**
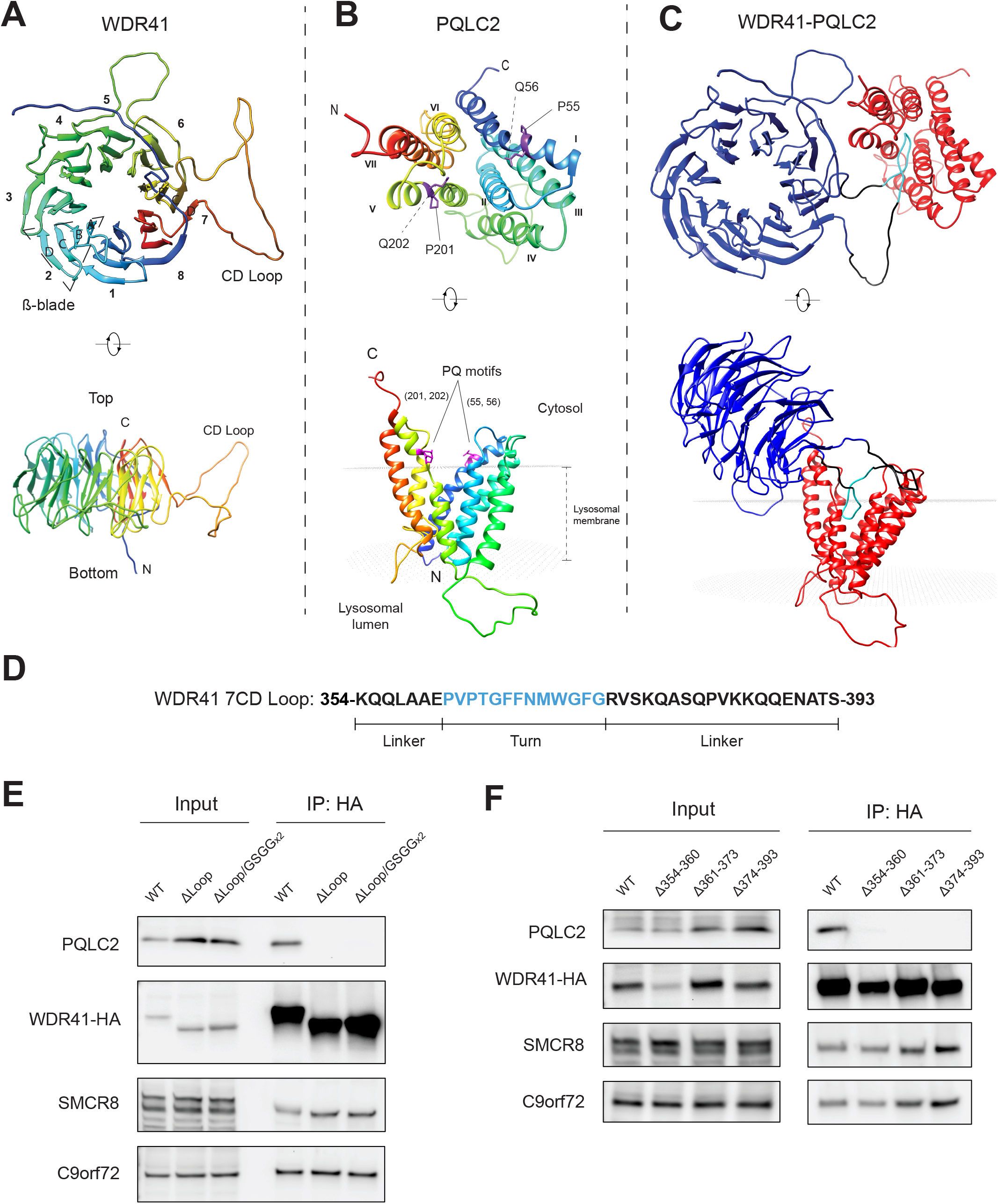
Generation and validation of a structural model predicting that the WDR41 7CD loop is necessary for interaction with PQLC2. (A) Ribbon diagram of the predicted WDR41 structure which forms an 8-blade β-propeller with a large loop (CD Loop) formed between the C and D β-strands in blade 7. (B) Predicted structure of PQLC2 inward facing conformation. PQ motifs in transmembrane helices 1 and 5 are highlighted. The position of PQLC2 in the lysosomal membrane was predicted via the Positioning of Proteins in Membranes (PPM 2.0) server (45). (C) Model derived from the docking of predicted WDR41 and PQLC2 structures (highest score model shown) wherein the 7CD loop of WDR41 is inserted within the central cavity exposed by the inward facing conformation of PQLC2. (D) WDR41 7CD loop amino acid sequence. (E) Anti-HA immunoprecipitation (IP) from cells expressing PQLC2-FLAG and HA-tagged wild-type or indicated mutant versions of WDR41-HA followed by detection of the indicated proteins by immunoblotting. (F) Anti-HA immunoprecipitation followed by immunoblot detection of HeLa cells expressing either WT WDR41-HA or WDR41-HA mutants with deletions to the regions depicted in 1D (linker, turn and linker) and PQLC2-FLAG.

In parallel, we predicted a structure for PQLC2 based on homology to another member of the PQ-loop transporter family known as SWEET13 from *Arabidopsis thaliana* whose structure (PDB: 5XPD) was previously solved in the inward facing conformation (Han et al., 2017). The resulting model for PQLC2 matches expectations for members of this family in that it contains an N-terminal three helix bundle (THB) made up by transmembrane domains (TMDs) 1-3 linked to the C-terminal THB made up of TMDs 5-7 via the TMD 4 linker [Fig. 1B; (25–27). This conformation contains a central cavity oriented towards the cytoplasm. The PQ-motifs in TMDs 1 and 5 were predicted to be located at the membrane-cytosol interface [Fig. 1B, side-view; inferred with the assistance of PPM web server (Lomize et al., 2012)].

The predicted structures of WDR41 and PQLC2 were used to perform ab initio asymmetric protein-protein docking via GalaxyTongDock (Park et al., 2019). In the highest ranked model emerging from these predictions the interaction was mediated by insertion of the 7CD loop of WDR41 into the central cavity of PQLC2 (Fig. 1C). In this model, the tip of the WDR41 7CD loop occupies a space within the central cavity close to where a substrate analogue was found to reside in the SWEET13 inward facing structure (Han et al., 2017).

### Mutagenesis experiments validate structural predictions

We next generated WDR41 mutants with either a complete deletion of the region encompassing the 7CD loop (residues 354-393) or its replacement by a flexible linker sequence (GGSGGGSG) and used the strong, constitutive, interaction between over-expressed WDR41 and PQLC2 to test our prediction about the WDR41-PQLC2 interaction mechanism. Immunoprecipitation (IP) experiments revealed that neither of the WDR41 loop deletion mutants bound to PQLC2. However, both mutants retained their ability to interact with C9orf72 and SMCR8 which argues that folding of their β-propeller remains intact (Fig. 1D). To identify determinants within the WDR41 loop for PQLC2 interaction we defined 3 sub-regions corresponding to the turn region which was predicted to insert most deeply into the PQLC2 inward facing cavity as well as the 2 flanking linker sequences (Fig. 1D) and tested the impact of deleting each of these sub-regions. Although none of these mutants interacted with PQLC2, they all retained interactions with C9orf72 and SMCR8 (Fig. 1E).

### A short motif within the WDR41 7CD loop is sufficient for PQLC2 binding

Based on the predicted model for WDR41-PQLC2 interactions (Fig. 1C), the negative impact of deletions to multiple parts of the WDR41 loop could reflect a need for specific sequences within the WDR41 turn region at the proposed site of contact with PQLC2 as well as flanking linker regions that extend the tip of this loop away from the β-propeller surface. To test this hypothesis, we fused the entire WDR41 loop sequence (354-393) or three small portions [(354-360 (linker); 361-373 (turn); and 374-393 (linker)] to the flexible tail extending from the C-terminus of EGFP (Fig. 2A). Anti-GFP immunoprecipitations revealed that the entire WDR41 7CD loop sequence (39 residues) as well as the GFP fusion protein containing the WDR41 turn region (361-PVPTGFFNMWGFG-373), but not either of the flanking sequences, interacted with PQLC2 (Fig. 2B). Further deletions to the turn region established that the 10 amino acids defined by WDR41 amino acids 364-373 (TGFFNMWGFG) represents the minimal linear peptide sequence able to interact with PQLC2 which we henceforth refer to as WDR41 PQLC2-interacting peptide (PIP, Fig. 2B). As a negative control, no interactions were observed between any of these GFP-fusions and SMCR8. Consistent with the critical role for the PIP sequence in mediating the interaction with PQLC2, multiple sequence alignments of the WDR41 7CD loop from diverse species revealed that the PIP region is more highly conserved than its flanking sequences (Fig. S1).

**Figure 2:**
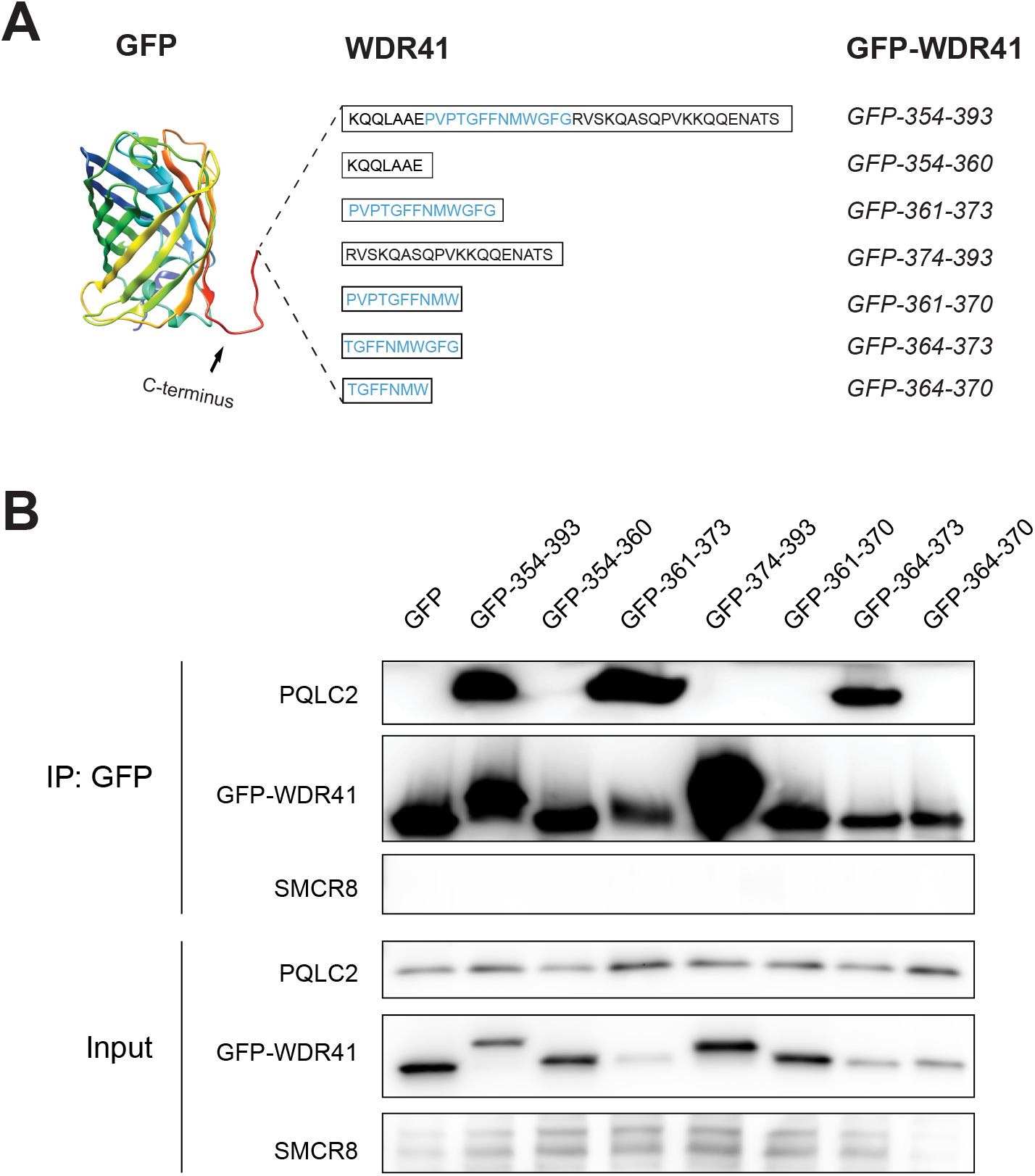
A 10 amino peptide from the WDR41 7CD loop is sufficient for PQLC2 interaction. (A) Schematic representation of GFP chimeric proteins containing pieces of the WDR41 7CD loop. The indicated amino acid sequence of WDR41 regions of interest were fused to the unstructured C-terminal extension of GFP. (B) Analysis of PQLC2-FLAG binding to GFP-WDR41 chimeras by immunoprecipitation and immunoblotting. Proteins were immunoprecipitated with GFP-Trap beads and samples corresponding to the lysates (Input) and immunoprecipitated proteins (IP: GFP) were immunoblotted with FLAG, GFP and SMCR8 antibodies.

A series of site-directed mutagenesis experiments were carried out to define key residues within the WDR41 PIP that are required for PQLC2 binding (Fig. 3A). Consistent with our previous experiments that defined the minimal PIP as amino acids 364-373, mutating prolines 361 and 363 to alanine had no impact on the interaction between full length WDR41 and PQLC2. Although the WDR41-G365A mutant still interacted with PQLC2, the bulkier G365V substitution at this site abolished the interaction. However, glycine to valine substitutions at 371 and 373 were both tolerated (Fig. 3B). Having observed the variable importance of specific amino acids within this WDR41 PIP region, we next performed an alanine scanning mutagenesis of this whole region which revealed that replacement of F366, F367, N368, M369, W370 or F372 with alanine abolished PQLC2 interactions (Fig. 3D). Meanwhile, the T364A (like G365A) mutation had minimal effects (Fig. 3D). Collectively, the selective importance of multiple amino acids within the WDR41-PIP sequence strengthens our conclusion that this peptide motif within the 7CD loop of WDR41 mediates the interaction with PQLC2. As all WDR41 mutants retained their interaction with SMCR8 (Fig. 3E), these changes within the 7CD loop did not grossly disrupt the β-propeller architecture of WDR41.

**Figure 3:**
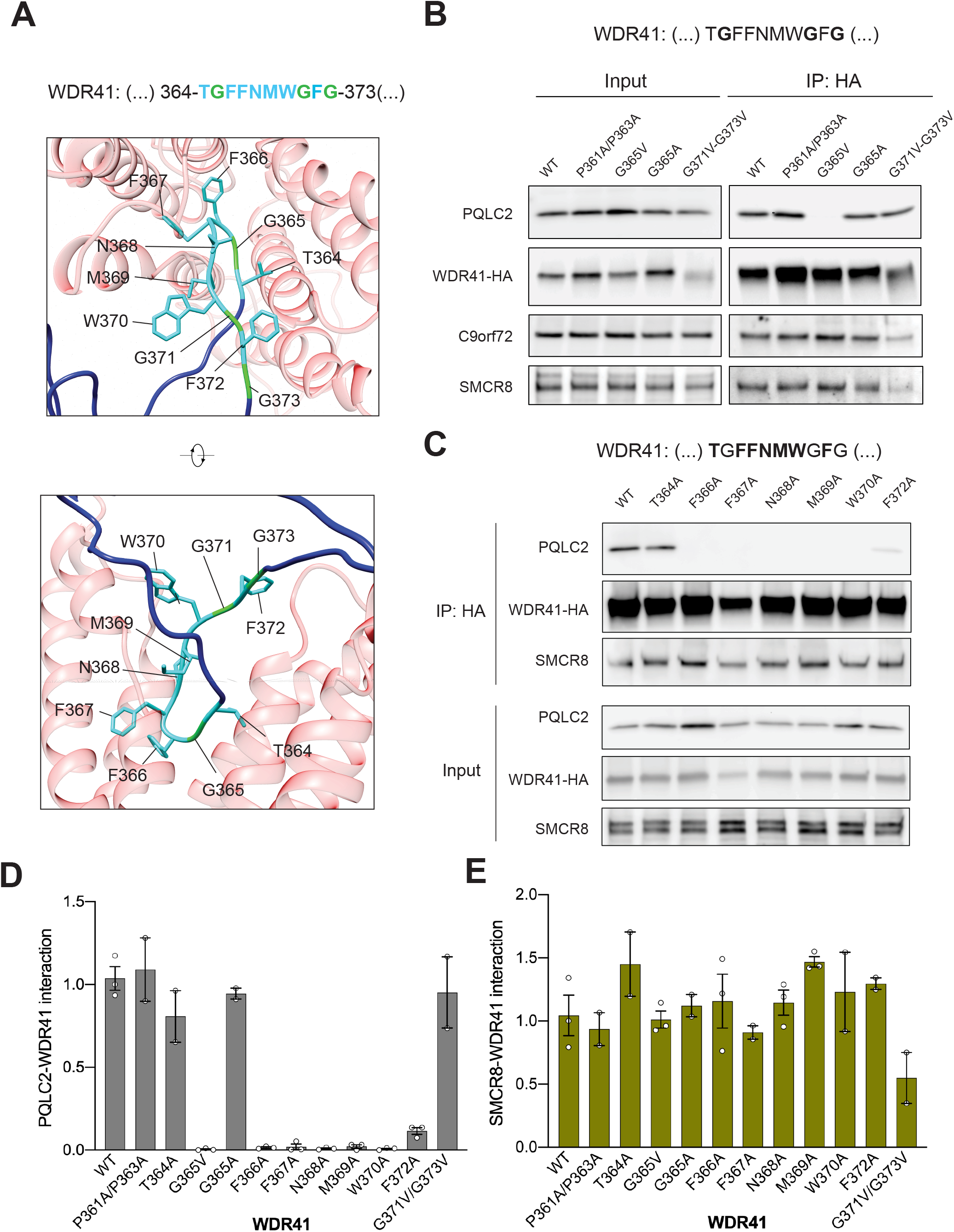
Site directed mutagenesis defines WDR41 specific residues in the 7CD loop that are required for PQLC2 binding. (A) Structural representation of the predicted interaction between WDR41-PQLC2 (frontview and sideview), showing the side-chains of the WDR41 PQLC2 interacting peptide (PIP) residues. (B) Impact of site directed mutagenesis of selected glycine and prolines within and adjacent to the WDR41-HA PIP on PQLC2-FLAG interactions. (C) Alanine-scanning mutagenesis of the remaining residues within WDR41 PIP (T364A, F366A, F367A, N368A, M369A, W370A or F372A). HeLa cells transiently co-transfected with plasmids expressing PQLC2-FLAG and WDR41-HA wildtype and mutants were lysed and immunoprecipitated (IP: HA) and immunoblotted with FLAG, HA, C9orf72 and SMCR8 antibodies. (D-E) Quantification of the PQLC2 and SMCR8 interaction with WDR41 mutants (normalized to wild-type WDR41). Mean ± SEM plotted with data points from independent experiments as open circles.

### WDR41 interacts directly with PQLC2 via the PIP motif

To test for a direct interaction between the WDR41 PIP and PQLC2, we performed pull-down assays with a recombinant protein consisting of GST fused to the 10 amino acid PIP sequence. When immobilized on beads, this fusion protein ineracted with PQLC2-FLAG from cell lysates (Fig. 4A). In reciprocal interaction experiments, where PQLC2-FLAG was first purified by immunoprecipitation and then used as bait, it selectively interacted with the recombinant GST-PIP fusion protein but not GST alone (Fig. 4B). These results corroborate the previous interaction assays using GFP-fusions and support a direct interaction between WDR41 PIP and PQLC2.

**Figure 4:**
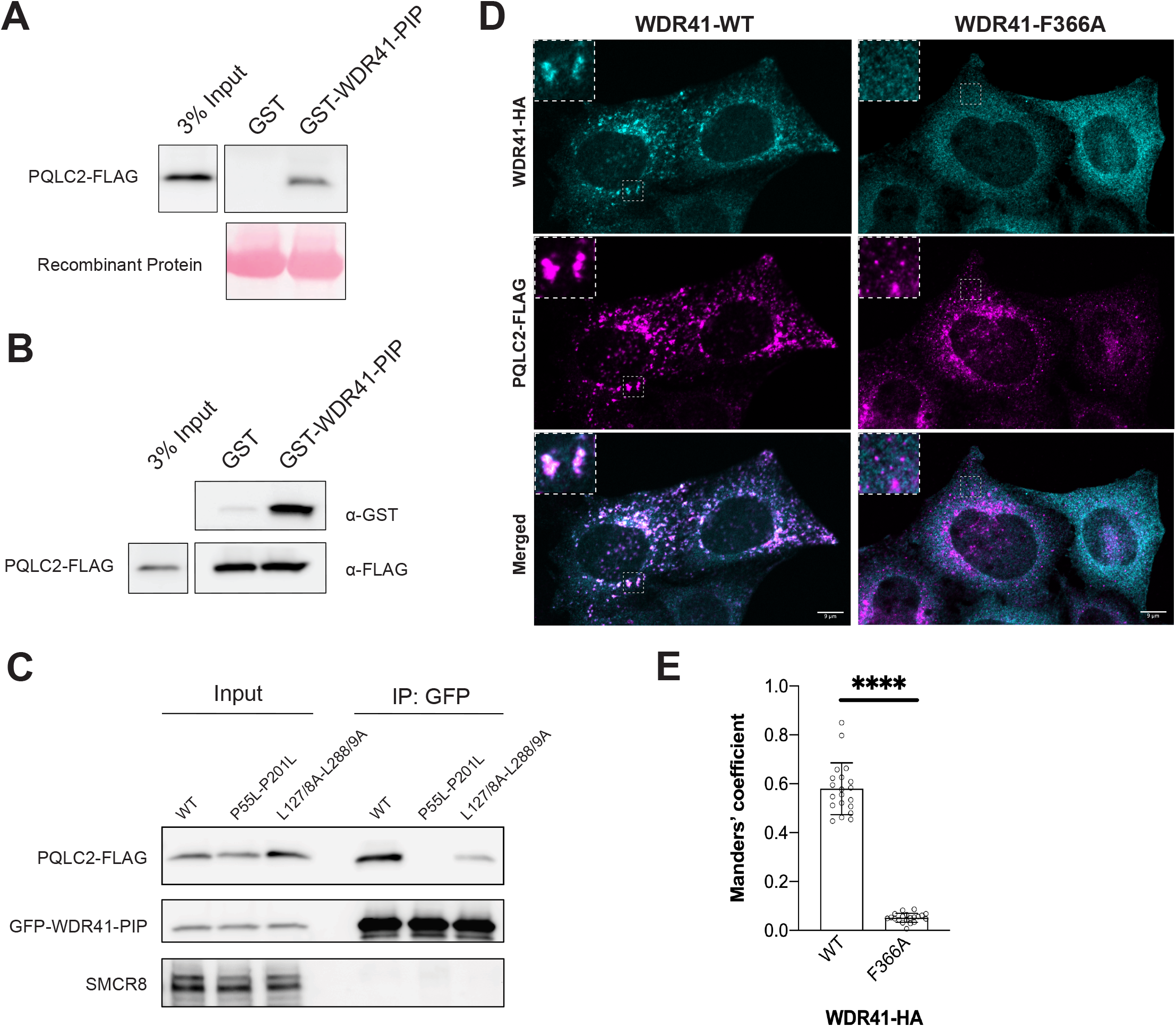
Direct interactions between WDR41 7CD loop and PQLC2 support WDR41 recruitment to lysosomes. (A) GST pull-down assay testing for the interaction between recombinant GST versus GST-WDR41-PIP in binding to PQLC2-FLAG. (B) PQLC2-FLAG was isolated by immunoprecipitation and used as bait to test for interaction with recombinant GST versus GST-WDR41 PIP. (C) Transiently co-transfected HeLa cells, expressing GFP-WDR41-PIP and FLAG-tagged PQLC2 wildtype or mutants (P55L-P201L or L127/8A-L288/9A), were lysed and immunoprecipitated with GFP-trap beads. Immunoprecipitated proteins (IP: GFP) and lysates (Input) were immunoblotted with FLAG, GFP and SMCR8 antibodies. (D) Spinning disc confocal immunofluorescence microscopy of transiently co-transfected HeLa WDR41 KO cells expressing PQLC2-FLAG and WDR41-HA-WT versus the WDR41-HA-F366A mutant. Insets represent a 3X magnification of the selected region, depicted by the dashed line box. Scale bar = 9 μm. (E) Quantification of WDR41 lysosome localization (PQLC2-FLAG colocalization) for WDR41-HA-WT versus WDR41-HA-F366A (Manders’ colocalization coefficient with threshold corrections where values represent the fraction of WDR41 overlapping with PQLC2). Mean ± SD is plotted with each dot representing the average of 10-20 transfected cells of a biological replicate (n=3), unpaired t test; p=0.0001; ***.

### PQLC2 conformational changes link its transporter and receptor-like functions

The PQ motifs in transmembrane helices 1 and 5 of PQLC2 are thought to functions as hinges that support conformational changes required for substrate transport and proline to leucine mutations at these sites block transporter activity of PQLC2 (22, 28, 29). In particular, this hinge is critical for the inward facing conformation of this family of transporters (30). Consistent with the predicted importance of the inward facing conformation for the WDR41 PIP interaction, the PQLC2 PQ motif proline to leucine mutants did not interact at all with the WDR41 PIP (presented as a GFP fusion protein; Fig. 4C). This result is further supported by our previous observation that although this PQLC2 mutant still localizes to lysosomes, it cannot recruit the WDR41-SMCR8-C9orf72 complex (5). Transporters that also have receptor-like signalling activities have been referred to as “transceptors” (31–33). Although the transceptor concept is supported by multiple pieces of functional data, mechanisms that relate the distinct transporter and receptor activities of transceptors have been elusive. Our findings which identify a conformational state of the alternating access transporter model in mediating WDR41 recruitment to the cytoplasmic surface of lysosomes thus provides a model for defining the basis for transceptor properties of PQLC2.

We also sought to test whether the subcellular localization of PQLC2 affects its ability to interact with WDR41. However, in contrast to previous studies where mutation of a single dileucine motif within the cytoplasmic C-terminus resulted in the accumulation of the rat PQLC2 on the plasma membrane rather than lysosomes (22, 34) we found that human PQLC2 contains two distinct dileucine motifs that must both be mutated to alanine in order to yield a plasma membrane rather than lysosome localization (Fig. S2). This plasma membrane-localized PQLC2 mutant exhibited reduced interaction with the WDR41-PIP GFP fusion protein (Fig. 4C). This may reflect a requirement for an acidic luminal pH and/or unique aspects of the lysosomal membrane for promoting the inward facing PQLC2 conformation that we predict to be a prerequisite for WDR41 interactions.

### WDR41-PIP is required for PQLC2-dependent recruitment of WDR41 to lysosomes

To test our model wherein the WDR41 loop mediates PQLC2 interactions and WDR41 recruitment to lysosomes in cells, we analysed subcellular localization of the WDR41-F366A mutant as a representative example of a PIP motif point mutant that cannot interact with PQLC2 (Fig. 3). Whereas, the wild-type WDR41 was enriched on PQLC2-positive lysosomes, the WDR41-F366A mutant was not recruited to lysosomes and was instead dispersed throughout the cytoplasm (Fig. 4D and E).

### A new foundation for understanding signalling from lysosomes in health and disease

We have provided a new model for the regulated, PQLC2-mediated, recruitment of the C9orf72 complex to the cytoplasmic surface of lysosomes based on the insertion of a short peptide motif within the WDR41 7CD loop region (WDR41-PIP) into the large cavity that is predicted to be formed by the inward facing conformation of PQLC2. In addition to the fundamental biological importance of understanding how this interaction allows cells to maintain lysosome homeostasis and adapt to changes in cationic amino acid availability (12, 15–19, 35). This raises the possibility that drugs which stabilize a conformation of PQLC2 that supports WDR41 interactions could have therapeutic value by enhancing C9orf72-SMCR8-WDR41 signalling from lysosomes.

## METHODS

### Structural predictions

Database searching and secondary structure prediction was performed with HHpred (https://toolkit.tuebingen.mpg.de/tools/hhpred), a sensitive protein homology detection and structure prediction by HMM-HMM comparison (36), core of an MPI Bioinformatics Toolkit (37). The modelling of WDR41 and PQLC2 was achieved with the Modeller, a program for comparative protein structure modelling (38) as part of the MPI Bioinformatics Toolkit (37). WDR41 secondary structure prediction was also accessed by WD40-repeat protein Structures Predictor (WDSP) (39). Protein-protein docking analysis was performed by GalaxyTongDock in an *ab initio* asymmetric fashion (40). Templates and models were analysed and selected based on HHpred and GalaxyTongDock criteria and scores. Visualization and 3D analysis were performed with UCSF Chimera (http://www.rbvi.ucsf.edu/chimera) (41).

### Cell Culture and Transfection

HeLa cells (provided by Pietro De Camilli, Yale University) were grown in Dulbecco’s Modified Eagle Medium (DMEM) + 4.5 g/L D-glucose, L-Glutamine (Gibco), 10% fetal bovine serum, and 1% penicillin/streptomycin supplement (Mediatech). Transfections and co-transfections were performed using FuGENE 6 transfection reagent (Promega). Cells were analysed two days post-transfection.

### Plasmids

PQLC2-FLAG plasmids were described previously (5). The WDR41-HA plasmid (pCMV6-WDR41-HA) was a gift from Nicolas Charlet-Berguerand (Addgene plasmid # 74159). Deletions and single/double substitutions of WDR41 gene were incorporated into WDR41-HA plasmid. GFP-WDR41 fusions were constructed in the pEGFP-N1 plasmid (Clontech) and the GST-WDR41 fusion was generated in pGEX-6T-1 (GE Life Sciences). These new plasmids were generated by Gibson Assembly or Q5 Site-Directed Mutagenesis according to manufacturer instructions (New England Biolabs). Primers for generating these mutants are listed in **Table S1**. Plasmids were confirmed by restriction enzyme digestion and sequencing (Yale University Keck DNA Sequencing Facility).

### Immunoprecipitation and immunoblotting

Cells were washed with cold PBS and then lysed in 50 mM Tris pH 7.4, 150 mM NaCl, 1mM EDTA, 1% Triton X-100 plus protease and phosphatase inhibitor cocktail tablets (Complete mini, EDTA-free, Phos-STOP from Roche). Insoluble cell material was rejected by centrifugation. Immunoprecipitations were performed on the resulting lysates. Anti-HA Affinity Matrix (Roche Diagnostics), GFP-Trap (ChromoTek, Planegg-Martinsried, Germany) and anti-FLAG M2 affinity gel (Sigma-Aldrich) were used for anti-HA, Anti-GFP and anti-FLAG immunoprecipitations, respectively. Samples were resolved in 4–15% gradient Mini-PROTEAN TGX precast polyacrylamide gels (Bio-Rad) and transferred to nitrocellulose membranes (Bio-Rad). Membranes were then blocked with 5% non-fat dry milk for 1h, and antibodies were incubated with 5% non-fat dry milk or bovine serum albumin (BSA), in TBS with 0.05% Tween 20, overnight. The antibodies used for western blot analysis were HA (3F10, Roche, 1:2000), GFP-HRP (Rockland, 1:2000), FLAG (Cell Signaling Technology, 1:1000), C9orf72 (GT1553, Genetech, 1:2000), SMCR8 (Bethyl Laboratories, 1:4000) and GST (Rabbit anti-GST antibody raised against full-length recombinant *Schistosoma japonicum* GST, provided by P. De Camilli, 1:2000). Chemiluminescence detection of horseradish peroxidase signals from primary or secondary antibodies was acquired by Versa-Doc Imaging (Bio-Rad) and FIJI (Schindelin et al., 2012) was used to quantify band intensities.

### GST fusion protein purification and pull-down assays

Plasmids for expression of GST fusion proteins were transformed in BL21 *E. coli*. The GST-tagged proteins were expressed and purified using Glutathione Sepharose 4B GST-tagged protein purification resin (GE Healthcare) following previously published protocols (42). Dialysis was performed via 3.5k MWCO Slide-A-Lyzer-G2 Dialysis Cassettes (ThermoScientific). Pull down assays were performed by adding GST-tagged proteins to the lysates from HeLa cells, transiently expressing PQLC2-FLAG plasmids, or HEK293FT C9orf72-HA PQLC2 KO cells (5), stably expressing PQLC2-FLAG (pLVX-puro vector, Clontech). Proteins bound to glutathione resin were then separated by SDS-PAGE and immunoblotted with anti-FLAG antibodies while the GST-tagged proteins were visualized by Ponceau S staining (APACOR). Inversely, FLAG immunoprecipitation in the presence of GST and the peptides, detecting GST-tagged proteins with GST antibody.

### Immunofluorescence and microscopy

HeLa WDR41 knockout cells (Amick et al., 2018) grown on 12 mm coverslips (Carolina Biological Supply) were transiently transfected with PQLC2-FLAG and WDR41-HA or WDR41-F366A-HA plasmids. Cells were fixed by a 1:1 addition of 8% paraformaldehyde (PFA) in 0.1 M sodium phosphate to their growth media, blocked in TBS with 5% NDS (Normal Donkey Serum) and antibody incubations were performed in the same buffer (Amick et al., 2020; Petit et al., 2013). Antibodies used in this experiment are described in Supplemental Table 2. A Nikon Inverted Eclipse TI-E Microscope equipped with 60x CFI PlanApo VC, NA 1.4, oil immersion and 40x CFI Plan Apo, NA 1.0, oil immersion objectives, a spinning disk confocal scan head (CSU-X1, Yokogawa) and Volocity (PerkinElmer) software was used for spinning-disk confocal microscopy. Manders’ Coefficients (with threshold corrections, M1: fraction of WDR41-HA or mutant overlapping PQLC2-FLAG) were calculated with JACoP colocalization plugin using FIJI (43, 44).

## ACKNOWLEDGEMENTS

We are grateful to all members of the Ferguson lab for helpful advice and thoughtful discussions. This research was supported by NIH grant GM105718 to SMF. The authors have no conflicts of interest to declare.

