## Supplemental Material for "Receptor-like role for PQLC2 amino acid transporter in the lysosomal sensing of cationic amino acids"

Shawn M. Ferguson

**This PDF file includes:**

Figures S1 to S2

Table S1

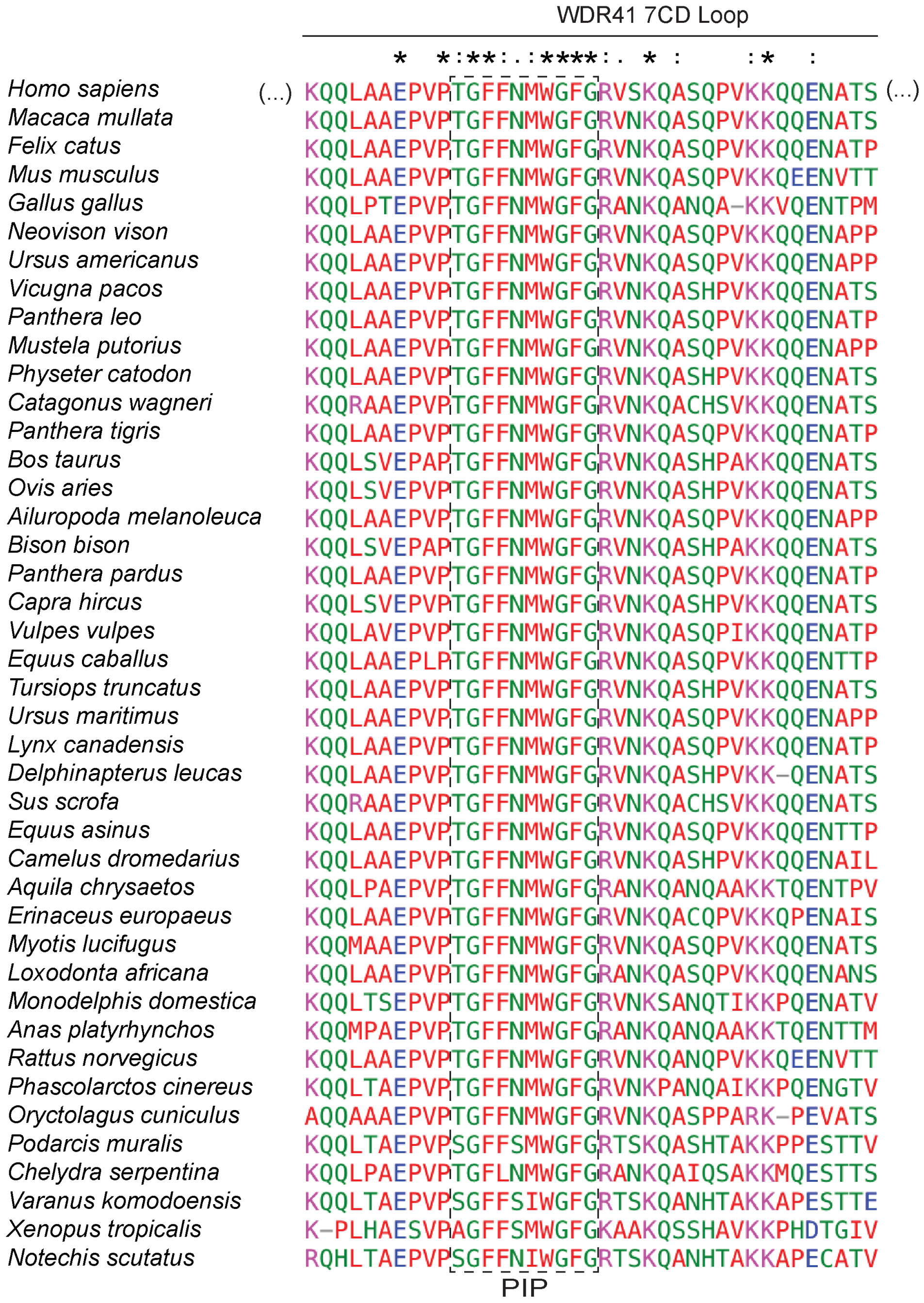


**Fig. S1. Selected amino acids are highly conserved within vertebrate WDR41 7CD loops.** Multiple sequence alignments of the WDR41 7CD loop from mammals, birds, reptiles and amphibians reveal a highly conserved peptide within the Turn region (defined in Figure 1) and which we subsequently defined as the PQLC2 Interacting Peptide (PIP). Protein sequences were obtained in Ensembl database (EMBL-EBI) and multiple alignment, using the entire sequence, was performed by Clustal Omega (EMBL-EBI). "*" fully conserved residue; "**:**" conservation between groups of strongly similar properties; "**.**" conservation between groups of weakly similar properties.

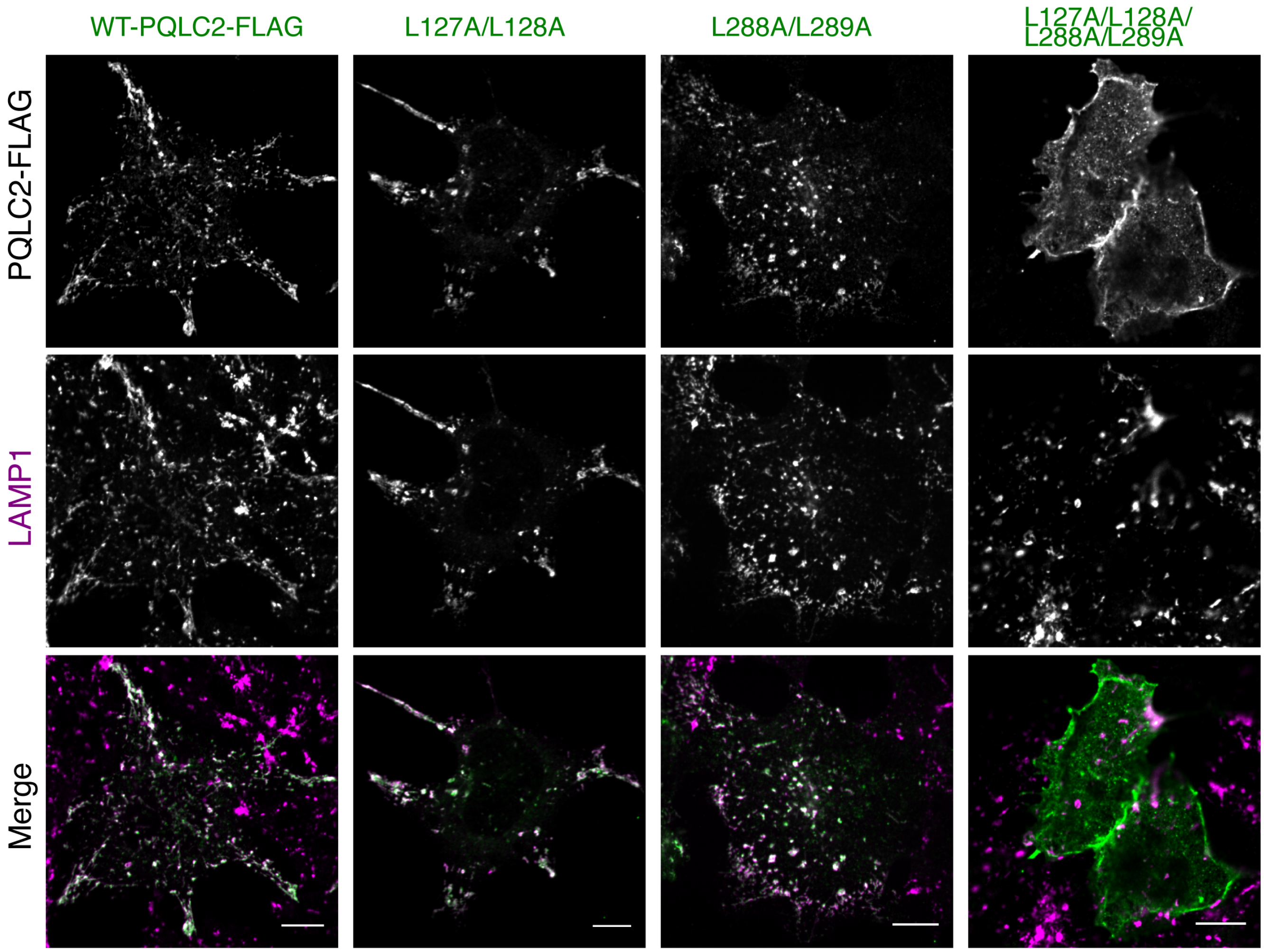


**Fig. S2. Identification of dileucine motifs required for lysosome localization of PQLC2.** Spinning disc confocal immunofluorescence microscopy of HEK293FT cells expressing PQLC2-FLAG with the indicated mutations along with detection of endogenous LAMP1 (late endosomes and lysosomes). Scale bar: 10μm.

### Table S1. Summary of oligonucleotide primers used in the study

| Gene/ plasmid | Use | Sense Sequence (5'-3') | AntiSense Sequence (5'-3') | Result |
|---|---|---|---|---|
| pCMV6-WDR41-HA | Deletion of 39 residues (Loop, 354-393) | TCACTGGAGCTTATTGGAG | TTCTCTTAACTCCCAAATGC | WDR41ΔLoop |
| pCMV6-WDR41-HA | Substitution of 39 residues (Loop) by (GSGG)x2 | GGCGGCAGCGGCTCACTGGAGCTTATTGGAG | GCCGCTGCCGCCTTCTCTTAACTCCCAAATGC | WDR41ΔLoop/GSGGx2 |
| pCMV6-WDR41-HA | Deletion of 354-360 | CCTGTACCAACAGGTTTTTTTAAC | TTCTCTTAACTCCCAAATGC | WDR41Δ354-360 |
| pCMV6-WDR41-HA | Deletion of 361-373 | AGAGTCAGCAAACAAGCC | CTCAGCTGCAAGCTGCTG | WDR41Δ361-373 |
| pCMV6-WDR41-HA | Deletion of 374-394 | TCACTGGAGCTTATTGGAG | TCCAAATCCCCACATGTTA | WDR41Δ374-393 |
| WDR41 | PCR (for DNA assembly) | GCATGGACGAGCTGTACAAGAAACAGCAGCTT GCAGCTG | ACATGAAGTAGCATTTTCTTGCTG | DNA fragment of WDR41LOOP (354-393) |
| pEGFP-N1 | PCR (for DNA assembly) | AAGAAAATGCTACTTCATGTTAAAGCGGCCGCG ACTCTAG | CTTGTACAGCTCGTCCATGC | Linearized plasmid |
| pEGFP-N1 | Insertion of 7 residues of WDR41 | TGCAGCTGAGTAAAGCGGCCGCGACTCT | AGCTGCTGTTTCTTGTACAGCTCGTCCATGC | GFP-354-360 |
| pEGFP-N1 | Insertion of 13 residues of WDR41 | TAACATGTGGGGATTTGGATAAAGCGGCCGCG ACTCT | AAAAAACCTGTTGGTACAGGCTTGTACAGCTCG TCCATGC | GFP-361-373 |
| pEGFP-N1 | Insertion of 20 residues of WDR41 | AAAAAGCAGCAAGAAAATGCTACTTCATGTTAA AGCGGCCGCGACTCT | AACAGGTTGGCTGGCTTGTTTGCTGACTCTCTT GTACAGCTCGTCCATGC | GFP-374-393 |
| pEGFP-N1 | Insertion of 10 residues of WDR41 | TAACATGTGGTAAAGCGGCCGCGACTCT | AAAAAACCTGTTGGTACAGGCTTGTACAGCTCG TCCATGC | GFP-361-370 |
| pEGFP-N1 | Insertion of 10 residues of WDR41 | TAACATGTGGGGATTTGGATAAAGCGGCCGCG ACTCT | AAAAAACCTGTCTTGTACAGCTCGTCCATGC | GFP-364-373 |
| pEGFP-N1 | Insertion of 7 residues of WDR41 | TAACATGTGGTAAAGCGGCCGCGACTCT | AAAAAACCTGTCTTGTACAGCTCGTCCATGC | GFP-364-370 |
| pCMV6-WDR41-HA | Double substitution | TAGCAACAGGTTTTTTTAACATGTGGG | CAGCCTCAGCTGCAAGCTGCTG | WDR41-P361A-P363A |
| pCMV6-WDR41-HA | Single substitution | GTACCAACAGTTTTTTTTAACATGTGGGG | AGGCTCAGCTGCAAGCTG | WDR41-G365V |
| pCMV6-WDR41-HA | Single substitution \ | GTACCAACAGCTTTTTTTAACATGTGGGGATTT G | AGGCTCAGCTGCAAGCTG | WDR41-G365A |
| pCMV6-WDR41-HA | Double substitution | TGTAAGAGTCAGCAAACAAGC | AATACCCACATGTTAAAAAAACC | WDR41-G371V-G373V |
| pCMV6-WDR41-HA | Single substitution | GCCTGTACCAGCAGGTTTTTTTAAC | TCAGCTGCAAGCTGCTGT | WDR41-T364A |
| pCMV6-WDR41-HA | Single substitution | ACCAACAGGTGCTTTTAACATGTGGGGATTTG | ACAGGCTCAGCTGCAAGC | WDR41-F366A |
| pCMV6-WDR41-HA | Single substitution | AACAGGTTTTGCTAACATGTGGGGATTTGGAAG | GGTACAGGCTCAGCTGCA | WDR41-F367A |
| pCMV6-WDR41-HA | Single substitution | AGGTTTTTTTGCCATGTGGGGATTTGGAAG | GTTGGTACAGGCTCAGCT | WDR41-N368A |
| pCMV6-WDR41-HA | Single substitution | TTTTTTTAACGCGTGGGGATTTGGAAGAGTC | CCTGTTGGTACAGGCTCA | WDR41-M369A |
| pCMV6-WDR41-HA | Single substitution | TTTTAACATGGCGGGATTTGGAAGAGTC | AAACCTGTTGGTACAGGC | WDR41-W370A |
| pCMV6-WDR41-HA | Single substitution | CATGTGGGGAGCTGGAAGAGTC | TTAAAAAAACCTGTTGGTAC | WDR41-F373A |
| pGEX-6T-1 | Insertion of 10 residues of WDR41 (PIP) | ATGTGGGGATTTGGAGGGATCCCCGGAATTCC C | GTTAAAAAAACCTGTGGGCCCCTGGAACAGAA C | GST-WDR41 PIP |
